# Detecting archaic introgression without archaic reference genomes

**DOI:** 10.1101/283606

**Authors:** Laurits Skov, Ruoyun Hui, Asger Hobolth, Aylwyn Scally, Mikkel Heide Schierup, Richard Durbin

## Abstract

Human populations out of Africa have experienced at least two bouts of introgression from archaic humans, from Neanderthals and Denisovans. In Papuans there is prior evidence of both these introgressions. Here we present a new approach to detect segments of individual genomes of archaic origin without using an archaic reference genome. The approach is based on a hidden Markov model that identifies genomic regions with a high density of single nucleotide variants (SNVs) not seen in unadmixed populations. We show using simulations that this provides a powerful approach to identifying segments of archaic introgression with a small rate of false detection. Furthermore our approach is able to accurately infer admixture proportions and divergence time of human and archaic populations.

We apply the model to detect archaic introgression in 89 Papuans and show how the identified segments can be assigned to likely Neanderthal or Denisovan origin. We report more Denisovan admixture than previous studies and directly find a shift in size distribution of fragments of Neanderthal and Denisovan origin that is compatible with a difference in admixture time. Furthermore, we identify small amounts of Denisova ancestry in West Eurasians, South East Asians and South Asians.

## Introduction

Archaic introgression into modern humans occurred at least twice (Neanderthals and Denisovans) (Meyer *et al*. 2012; Prufer *et al*. 2014) and had a phenotypic effect on humans (Huerta-Sanchez *et al*. 2014; Dannemann *et al*. 2017; Racimo *et al*. 2017). A substantial amount of Neanderthal and Denisovan genetic material is still present in modern humans and we can learn about archaic populations from studying their genetic variants in humans. To harness this information a number of methods have been developed to infer segments of archaic ancestry in an individual’s genome. Scanning along the genome, Hidden Markov Models (HMMs)(Prufer *et al*. 2014; Seguin-Orlando *et al*. 2014) and Conditional Random Fields (CRF)(Sankararaman *et al*. 2016) can identify haplotype segments in non-Africans that are both closer to the archaic reference genomes than to Africans, and also longer than expected by incomplete lineage sorting; these are then identified as likely archaic introgressed segments. Another approach is to identify segments with more variants in high linkage disequilibrium (LD) that are unique to non-Africans than expected given a certain demographic scenario (Plagnol and WALL 2006). The latest implementations of this method also use an archaic reference genome for refining the set of putative archaic haplotypes (Vernot *et al*. 2016).

The use of archaic reference genomes for identification of introgressed fragments has drawbacks. First, since the Neanderthal reference genomes are closer to the introgressing Neanderthal (80,000-145,000 years divergence)(Prufer *et al*. 2017), than the introgressing Denisova is to the Denisova genome (276,000-403,000 years divergence)(Prufer *et al*. 2014) detecting Denisovan ancestry will be harder. Second, the reliance on having reference genomes implies that the introgression maps generated by these methods need updates whenever more archaic reference genomes are sequenced (Prufer *et al*. 2017). Finally, it may be hard to identify introgressing segments of unknown archaic origin if such exists, as in the case of the putative archaic introgression into Pygmies (Hsieh *et al*. 2016) and Andamanese islanders (Mondal *et al*. 2016).

Here we present a new method for the identification of archaic introgression that does not require a reference genome or prior knowledge of demographic parameters, but uses density of variants in individuals private to their population of origin. We demonstrate with Papuans how we can estimate demographic parameters relevant to introgression and infer more archaic material than previously. Furthermore we can separate introgression events into Denisovan and Neanderthal components that display different length distributions in accordance with different admixture times.

## Method

### Model

An archaic genomic segment introgressed into a population is expected to have a high density of variants not found in populations without the introgression. We use a Hidden Markov Model (HMM) to classify genomic segments into states with varying density of such variants. We focus on a scenario where introgression with a deeply divergent archaic population only happened into an ingroup and not the outgroup, see Figure 1a. By removing variants found in the outgroup we can better distinguish introgressed segments from non-introgressed segments based on the density of remaining variants, see Figure 1a. These remaining variants, which we denote private variants (because they are private to the ingroup with respect to the outgroup) can either have occurred on the branch starting from the split of the ingroup and outgroup, or on the introgressing population’s branch. Because the introgressing segments have had a longer time to accumulate variants, they should have a higher density of private variants. Thus, we define a HMM with two states. The hidden states are Ingroup and Archaic, and the probability for changing state in the Ingroup is *p* and the probability for changing state in the Archaic is *q*, see Figure 1b. The probability of changing state can also be expressed in terms of a constant recombination rate between windows *r* · *L*, the admixture time *T*_*admix*_ and admixture proportion *a*, see Figure 1b.

**Figure 1.**
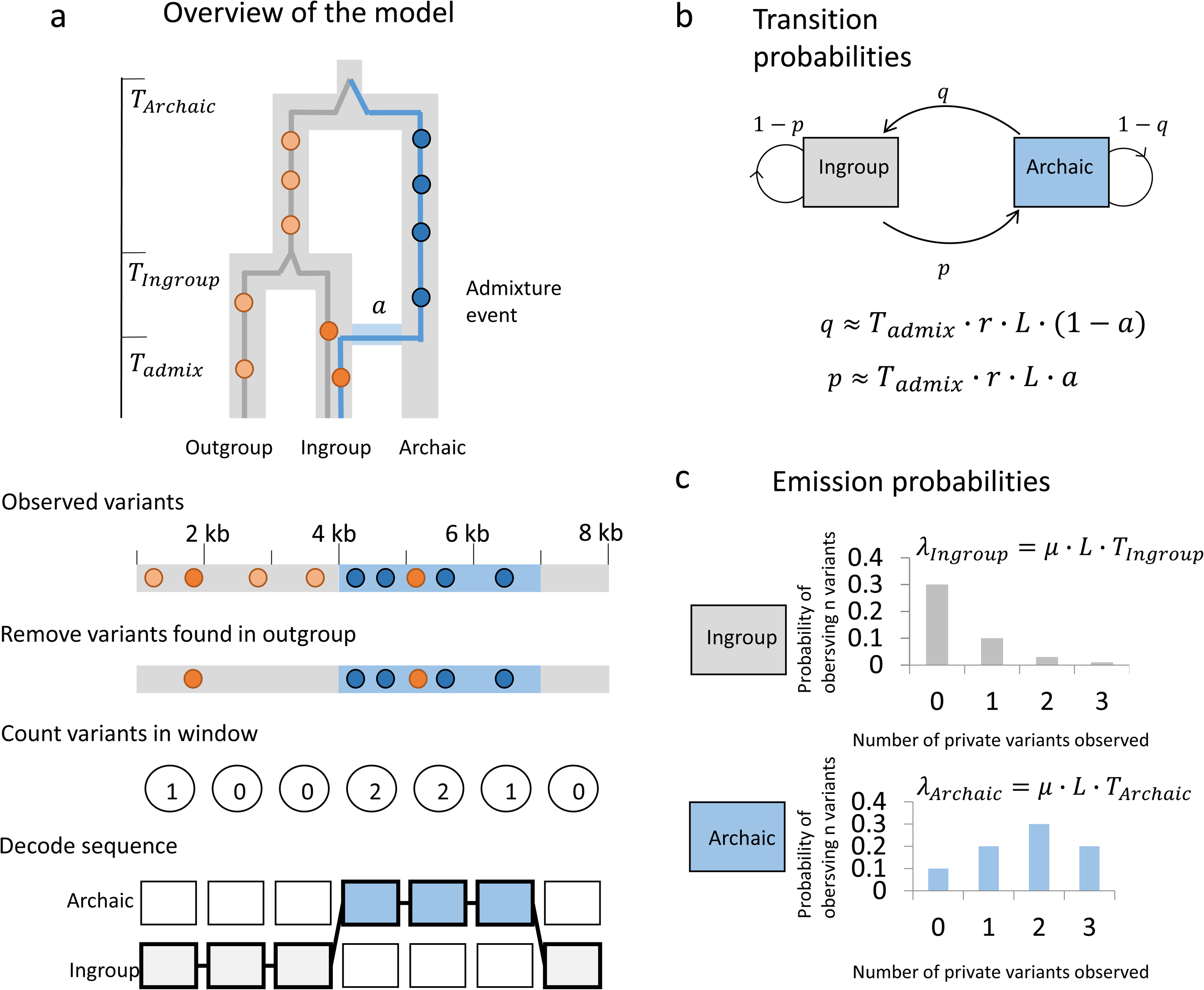
Overview of the model. Illustration on small test dataset. a) An archaic segment *T*_*admix*_ introgresses into the ingroup population at time T with admixture proportion a. The segments in the ingroup have a mean coalescence time with a segment from the outgroup at time *T*_*Ingroup*_ and an archaic segment has a mean coalescence time with a segment from the outgroup at time *T*_*Archaic*_. Removing all variants found in the outgroup (light orange points) should remove all the variants in the common ancestor of ingroup and outgroup, leaving only private variants that either occurred on the ingroup branch (dark orange) or on the archaic branch (dark blue). This will make the archaic segment have a higher variant density. The genome is then binned into windows of L (here 1000 bp) and the number of private variants are counted in each window. These are the observations and the hidden states are either Ingroup state or Archaic state. When decoding the sequence the most likely path through the sequence is found. b) The transition matrix between the archaic state and ingroup state. c) The emission probabilities are modelled as Poisson distributions with means *λ*_*Ingroup*_ and *λ*_*Archaic*_. It is more likely to see more private variants in the Archaic state than in the Ingroup state.

For practical purposes we bin the genome into windows of length L (typically *L = 1000 bp*). The number of private variants observed in a window is Poisson distributed with a rate *λ*_*Ingroup*_ and *λ*_*Archaic*_, respectively where *λ*_*Ingroup*_ *= µ* · *L* · *T*_*Ingroup*_ and *λ*_*Archaic*_ *= µ* · *L* · *T*_*Archaic*_, *µ* is the mutation rate, *T*_*Ingroup*_ is the mean coalescence time for the ingroup and the outgroup and *T*_*archaic*_ is the mean coalescence time for the archaic population and the outgroup, see Figure 1c.

We make a correction to the rates to take into account the number of missing bases in a window and the local mutation rate. For window *i* we have *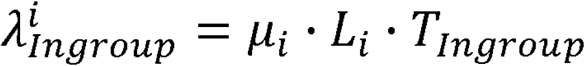* and *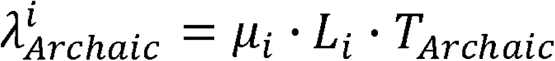* where *µ*_*i*_ is the local mutation rate and *L*_*i*_ is the number of called bases in a widow. The set of transition parameters *p, q* and the Poisson parameters *λ*_*Ingroup*_, *λ*_*Archaic*_ that maximize the likelihood given the observations are found using the Baum-Welch algorithm for an individual genome. These parameters are informative of the mean coalescence times between the ingroup and outgroup and between the archaic and the outgroup, the admixture time and the admixture proportion if we assume a known mutation rate *µ* and a known recombination rate between windows *rL*. Once the set of optimal parameters are found they can be used to decode the genome, using posterior decoding to identify candidate introgressed segments as consecutive regions with posterior probability of coming from the archaic state above some threshold.

To avoid problems with phasing we run this model on unphased diploid genomes. Heterozygous archaic segments will still stand out from homozygous non-introgressed segments. Formally this is equivalent to assuming that homozygous introgressed segments are sufficiently rare that they can be ignored for model fitting. In practice any homozygous archaic segments will have higher private variant density than heterozygous segments, so in the absence of a homozygous HMM state they will be classified with the heterozygous state.

## Results

### Testing the model with simulations

To investigate the ability of our model to identify archaic (Neanderthal and Denisovan) admixture into Papuans we simulated whole autosome diploid data using a coalescent simulator, with admixture with an archaic hominin 1,500 generations ago replacing 5% of the population – (a script with all demographic parameters are shown in Supporting information – Simulation script.py and a graphic representation of the demography is shown in Supporting figure 1). We simulated three scenarios to test the effects of missing data and varying recombination rate. The mutation rate was kept constant across the genome for all simulations.

First, we simulated five individuals where every base in the genome is called equally well and there is a constant recombination rate of 1.2 · 10^−8^ events per basepair per generation. We call this dataset the ideal data. Second, we simulated five individuals and removed all variants that are in repetitive regions (using the repeatmask track for the human reference genome hg19 (Smit *et al*. 2013)) to test how the model performs with missing data. Third, we simulated five individuals with missing data and using a varying recombination rate (using HapMap phase II (International Hapmap *et al*. 2007)) to test the effect of missing data and recombination. We binned all genomes into bins of 1000 bp, and removed all variants found in 500 simulated Africans, 100 simulated Europeans and 100 simulated Asians. We combine two haplotypes to form genotype data for the simulated individuals. This will be more similar to situations where phased data is not available.

We found the transition and emission parameters that optimized the likelihood, using the Baum-Welch algorithm and used them to get an estimate for the admixture time *T*_*admix*_, the admixture proportion *a* and the mean coalescent times with the outgroup *T*_*Ingroup*_ and *T*_*Archaic*_ for the ingroup and archaic segments respectively, see Figure 2b.

**Figure 2.**
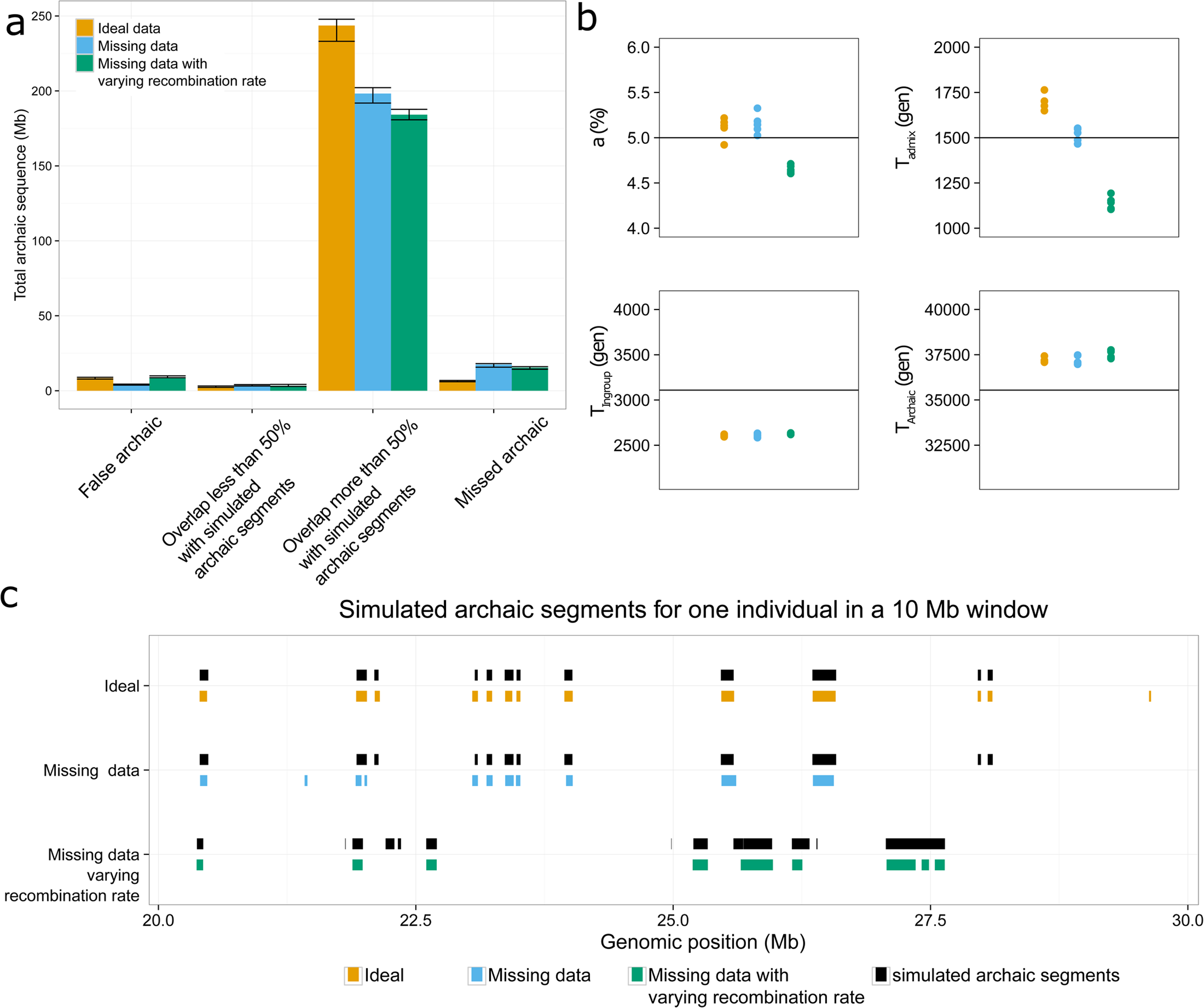
Evaluation of the model on simulated data. a) Average amount of sequence per individual that come from segments that are classified as false archaic (zero percent overlap with any true archaic segment), found < 50% (segment where there is less than 50 % overlap with true archaic segments), found > 50 % (segments where more than 50 % overlap with true archaic segments) and missed archaic which are segments that the model does not identify as archaic. The bars are colored according to what simulation scenarios they belong to. b) The estimation of the four parameters *T*_*admin*_, *a, T*_*Ingroup*_ and *T*_*archaic*_ are shown for the different simulation scenarios. c) An example of how simulated archaic segments and putative archaic segments overlap in a 10 Mb window. The x-axis is the genomic coordinates in Mb and the y-axis is the different simulation scenarios.

Across all scenarios the mean estimated coalescence time between the ingroup and outgroup (*T*_*Ingroup*_) is 2,625 generations (max = 2,647, min = 2,595), while the corresponding average simulated coalescent time was 3,109 generations ago. For the coalescence time between the outgroup and the archaic (*T*_*Archaic*_) the mean estimate is 37,345 generations (max = 37,832, min = 37,028) and the average simulated values was 35,543 generations ago.

We find that the mean estimate of the admixture proportion *a* when using the transition matrix is between 4.62 % and 5.34 %, consistent with the 5% simulation value.

We estimated the false negative rate of the model by counting the amount of simulated archaic segments that have zero overlap with the putative archaic sequence. This is 7.4 Mb for ideal simulations, 20.3 Mb for simulations with missing data and 17.5 Mb for simulations with missing data and a varying recombination rate, see Figure 2a. Most of this is in short segments which the model has less power to identify, as can be seen in Supporting figure 2.

If we estimate the false positive rate as the amount of inferred archaic segments that have zero overlap with the simulated archaic segments we find 16.6 Mb for ideal simulations, 13.2 Mb for simulations with missing data and 17.1 Mb for simulations with missing data and a varying recombination rate, see Figure 2a. We are therefore controlling for specificity (false positives) while losing sensitivity (false negatives) as the inference becomes more difficult. An example of how the simulated and putative archaic segments overlap is shown in Figure 2c for a 10 Mb segment.

We also find that with a posterior decoding threshold at 0.8 (mean posterior probability of being archaic for all windows in segment), the amount of false positives can be reduced by up to 50%, while still keeping 90% of the true segments, see Supporting figure 3. When applying a threshold of 0.8 we recover 246 Mb, 202 Mb and 187 Mb of archaic sequence for Ideal simulations, simulations with missing data and simulations with missing data and varying recombination rate respectively. When applying a threshold of 0.8 we recover 246 Mb, 202 Mb and 187 Mb of archaic sequence for Ideal simulations, simulations with missing data and simulations with missing data and varying recombination rate respectively.

**Figure 3.**
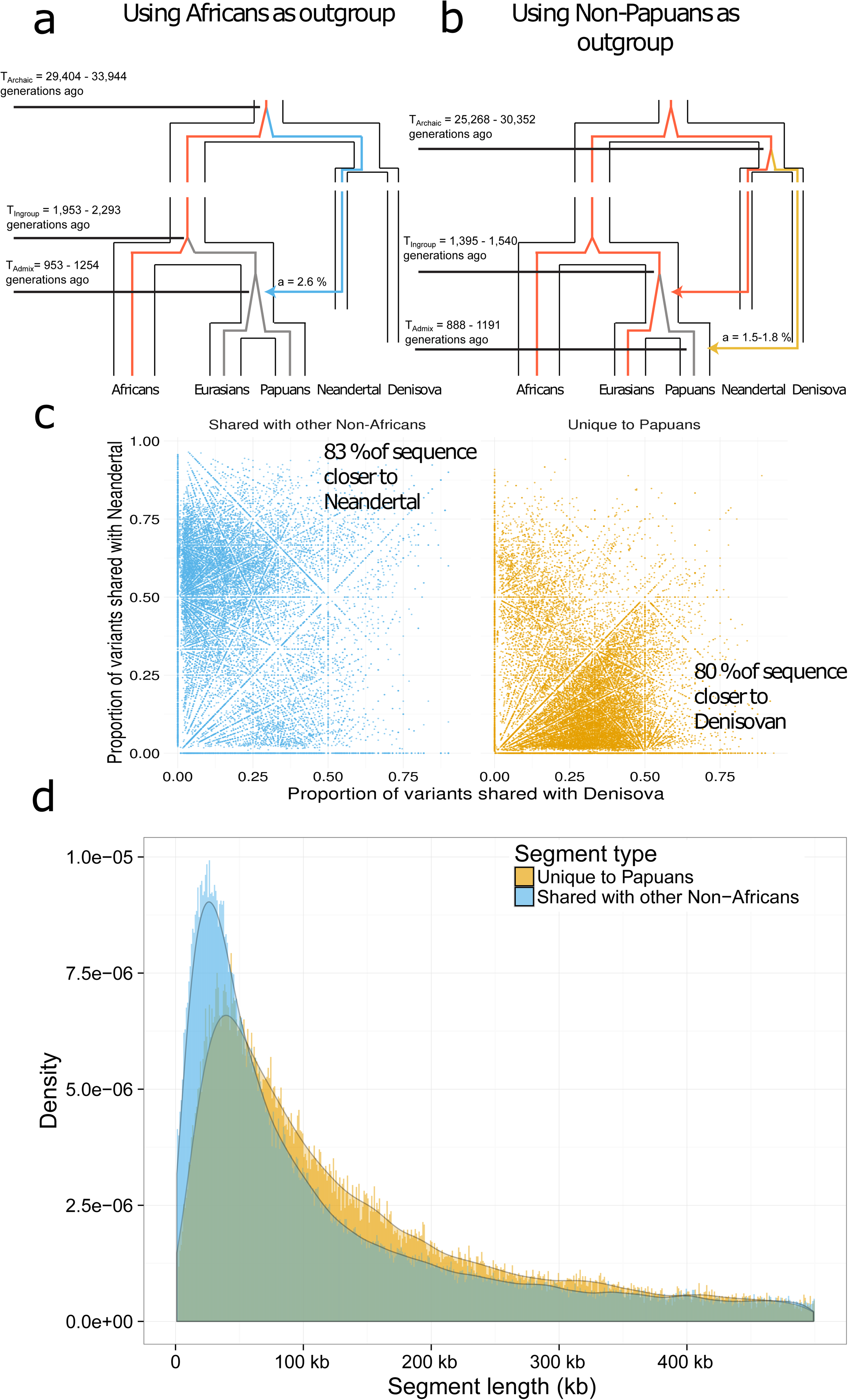
Application of model to Papuan genomes. a) Relationship between modern and archaic humans with the outgroup branches (Sub-Saharan Africans) colored in red. The average coalescence times for ingroup and outgroup *T*_*Ingroup*_ and archaic and outgroup *T*_*Archanic*_ are shown. The admixture proportions a and admixture time *T*_*admin*_ are shown for segments that are shared with other non-African populations. b) The outgroup colored in red is now all non-Papuans, and the new demographic parameters are shown. c) The segments that are shared with other Non-Africans share more variation with the Vindija Neanderthal than they do with the Altai Denisova. Segments that are unique to Papuan individuals share more variation with Altai Denisova than they do with the Vindijaarchaic segments with a mean posterior probability > 0.5 are kept) for segments that are shared with other non-African populations is shorter than segments that are unique to Papuans. segments with a mean posterior probability > 0.5 are kept) for segments that are shared with other non-African populations is shorter than segments that are unique to Papuans.

The mean estimate for the admixture time using the transition matrix is around 1,704 generations ago when using the ideal data and 1,522 generations ago when adding missing data. When we vary the recombination rate across the genome the average estimate of the admixture time is 1,146 generations ago if we estimate it using the transition matrix. The underestimate of the admixture time might be due to fact that the model fail to identify around 80% of the short segments in such cases. This would make the average segment length longer and make the admixture time seem more recent.

### Application to Papuan genomes

Having verified the validity of the model, we applied it to 14 Papuan individuals from the Simons Genome Diversity Project (Mallick *et al*. 2016), 40 Papuans from (Malaspinas *et al*. 2016) and an additional 35 Papuans (Vernot *et al*. 2016). In addition to this, we also analyzed individuals from West Eurasia, East Asia and South East Asia.

For each individual we used two different sets of variants as outgroup. We estimate the background mutation rate in windows of 100 kb, using the variant density of all variants in African populations from the 1000 Genomes Project.

Our model will not be able to distinguish Neanderthal from Denisova segments in Papuans, because the Denisovans and Neanderthals share a common ancestor before they do with humans and therefore the mean coalescence time with humans will be the same (Prufer *et al*. 2014). This means that the Poisson parameters will be the same as they both depend on *T*_*Archaic*_. However, we should be able to enrich for Denisova and Neanderthal segments by using different outgroups in our filtering step.

First, we used only variants found in Sub-Saharan African populations as an outgroup. This should remove variation in the common ancestor of Sub-Saharan Africans and the Papuans, retaining archaic variants of Neanderthal and Denisova origin as both are present in Papuans, but mainly absent in Africa (Sankararaman *et al*. 2016; Vernot *et al*. 2016). We also used this filter when analyzing Eurasian populations.

Second we remove variants found in all non Papuan populations, only retaining variants that are unique to Papuan populations. This should remove Neanderthal variants that are shared with other non-African populations (Prufer *et al*. 2014) and also to some extent remove variants of Denisovan origin that are found in Asians and Native Americans (Skoglund and Jakobsson 2011; Qin and Stoneking 2015). Thus removing all variants from the 1000 Genomes Project should enrich for Denisovan segments, while the segments that are found when using Sub-Saharan Africans but not using all 1000 Genomes Project samples as outgroups should be enriched for Neanderthal segments.

We found the optimal set of transition and emission parameters for each Papuan individual and found them to be largely consistent across the different datasets, see Supporting figure 4. The parameters were converted into estimates of *T*_*admix*_, a, *T*_*Ingroup*_ and *T*_*Archaic*_ using an average recombination rate of 1.2 · 10^−8^ events per basepair per generation and an average mutation rate of 1.25 · 10^−8^ mutations per base pair per generation, see Figure 3a, b.

We find that mean coalescence time between Papuans and non-Papuan individuals happened more recently (1,395-1,540 generations ago) than the mean coalescence time with Sub-Saharan Africans (1,953-2,293 generations ago) reflecting that Papuans are more closely related to other Non-Africans than to Africans. The mean coalescence time between Papuans and other non-Africans also provides an upper limit for Neanderthal introgression because it happened into the common ancestor of these populations.

Using only Sub-Saharan individuals as an outgroup we find a mean coalescence time between the archaic and outgroup to be between 29,404 and 33,944 generations ago. When using non-Papuans as an outgroup the estimate is between 25,268 and 30,352 generations ago. The lower estimate is likely due to the fact that some of the variants in the common ancestor of Denisovans and Neanderthals have been removed.

Using Sub-Saharan Africans as an outgroup we estimate the total admixture proportion of archaic sequence into Papuans to between 4.1-4.4 % and the admixture proportion that is private to Papuans between 1.5-1.8 %. This means that approximately 2.6 % is shared with non-Papuans, see Figure 3a.

From the transition parameters, we estimate that the admixture event with non-Africans happened 953-1,254 generations while the Papuan specific admixture event happened 888-1,191 generation ago. Both are likely underestimates as it was for the simulated data with missing data and varying recombination rate. Neanderthal admixture likely occurred closer to 2,000 generations ago after the out of Africa migration (Fu *et al*. 2014; Sankararaman *et al*. 2016) with Denisovan admixture occurring after that.

We used a threshold of 0.8 posterior probability as in the case of the simulated data. By comparing to the Vindija Neanderthal (Prufer *et al*. 2017) and Denisova (Meyer *et al*. 2012) genomes we find that this cutoff removes around 65% of the segments that don’t share variants with any archaic reference genome, see Supporting figure 5. These only contain 10.4 % of the total length of inferred archaic segments, and as well as including less confident segments may include deeply coalescing modern human haplotypes.

When we use a cutoff of 0.8 we find that 84 % of the segments unique to Papuans (80 % of the total sequence) shared more variants with the Denisova genome than with the Vindija Neanderthal, and that 78 % the segments that are shared with other non-Africans (83 % of the total sequence) shared more variants with the Vindija Neanderthal than the Denisova (Figure 3c). This is consistent with a majority of the archaic sequence unique to Papuans coming from a population more closely related to Denisovans, while a majority of the shared archaic sequence came from Neanderthals.

However, segments that are unique to Papuans are longer on average (94.2 kb) compared to those shared with other non-African populations (76.9 kb), See Figure 3d. The difference in length distributions are not seen as clearly when using Sstar or CRF, see Supporting figure 6. Moreover, the length distribution of archaic segments that are not unique to Papuans are more similar to other non-African populations, see Supporting figure 7.

We compared our archaic segments to those previously reported using other methods (Sankararaman *et al*. 2016; Vernot *et al*. 2016). We find that 67% of the archaic sequence found using CRF are also recovered using our method, and that 74% of the archaic sequence found using Sstar are also recovered using our method.

Comparing to the archaic reference genomes our method finds more Denisova in Papuans than it finds Neanderthal, unlike the CRF. It also finds a significant amount of additional Denisova segments in East and South East Asians, see Table 1.

**Table 1.** Amount of sequence of different origins. The amount of sequence (in Mb) that is equally related to Denisova and Vindija, more closely related to Denisova, doesn’t share any variation with either and is more closely related to Vindija are shown different populations and different methods.

| *Model* | *Pop* | *Both* | *Denisova* | *None* | *Vindija* | *Total* |
| --- | --- | --- | --- | --- | --- | --- |
| *HMM* | Papuan | 4.40 | 77.00 | 11.39 | 71.44 | 164.23 |
|  | eastasia | 1.48 | 5.69 | 9.96 | 61.37 | 78.49 |
|  | southasia | 1.62 | 5.85 | 10.12 | 51.36 | 68.95 |
|  | westeurasia | 1.47 | 2.39 | 10.14 | 43.95 | 57.94 |
| *Sstar* | Papuan | 26.5 | 43.11 |  | 49.21 | 118.82 |
|  | eastasia | - | 0.00 | - | 65.02 | 65.02 |
|  | southasia | - | 0.00 | - | 55.18 | 55.18 |
|  | westeurasia | - | 0.00 | - | 51.23 | 51.23 |
| *CRF* | Papuan | - | 58.17 | - | 84.72 | 142.89 |
|  | eastasia | - | 3.21 | - | 72.92 | 76.14 |
|  | southasia | - | 2.79 | - | 61.36 | 64.15 |
|  | westeurasia | - | 0.68 | - | 57.29 | 57.97 |

## Discussion

Since emission probabilities are very different between the human and archaic states in our model, we expect a low rate of false positive archaic inference, and this is also what we see in simulations. However, since recombination rates are highly variable, we expect many very short archaic segments and these have a very high false negative rate. Our inability to identify these causes us to underestimate the admixture time. However, the model does seem to find the correct size distribution for longer segments (> 50 kb), see Supporting figure 2. The mean coalescence times of modern and archaic humans are reasonably well estimated in simulations. One issue of interest is that the potential presence of super-archaic introgression as reported into the sequenced Denisovan (Prufer *et al*. 2014) should cause the mean coalescence time to Denisovan introgressed segments to be greater than that for Neanderthal segments. We did not observe this, perhaps because some Denisovan admixture is also present in East Asians who form part of our contrast population, reducing apparent mean divergence.

Our model reports more Denisova segments than approaches relying on the Denisovan reference. This is possibly because our method does not rely on putative Denisova segments being more closely related to the Denisova genome than the Vindija Neanderthal genome. Given that the introgressing “Denisovan” and the sequenced Denisova individual’s lineages split relatively shortly after the Neanderthals split from Denisovans (Prufer *et al*. 2014) many segments may be equally close to the Vindija Neanderthal and the sequenced Denisova sample. It is also expected that a fraction of segments introgressed from the Denisovan are more closely related to Vindija and vice versa due to incomplete lineage sorting. It is therefore also reassuring that we do not find the same large excess of Neanderthal fragments in Papuans compared to Asian populations as has been reported previously, see Table 1.

We find no clear evidence for an introgression with a new archaic hominin in Papuans, but we do find segments that do not share variation with any of the sequenced archaic populations. These segments could represent variation in Neanderthals and Denisovans that is not captured by the three high coverage archaic reference genomes, or another source. In the future it will be interesting to compare these segments to other human populations that might also have archaic segments of unknown origin (Hsieh *et al*. 2016; Mondal *et al*. 2016).

Our model is not restricted to being applied to humans. It works particularly well when it is possible to remove all the common variation between the ingroup and outgroup. As a larger number of individuals from different species are being sequenced, this method could be used as an alternative method for identifying introgression in other species, for example chimp and bonobo (De Manuel *et al*. 2016), bears (LIU *et al*. 2014), elephants (Palkopoulou *et al*. 2018) or gibbons (Carbone *et al*. 2014).

## Materials and methods

### Simulations

To simulate data we used Msprime (Kelleher *et al*. 2016). We simulated 5 Papuans and as an outgroup we simulated 500 Africans, 100 Europeans and 100 Asians using demographic parameters from (Malaspinas *et al*. 2016). We simulated data where we varied the recombination rate according to HapMap recombination maps (International Hapmap *et al*. 2007) for 5 individuals and removed variants within non-callable regions and variants that were found in the simulated outgroup. We grouped all autosomes into bins of 1000 base pairs and counted the number of variants. For each 1000 bp window we calculated the number of called bases using the repeat masked segments.

### Train parameters and decode segments

We trained and decoded the segments using our HMM, which is available at: https://github.com/LauritsSkov/Introgression-detection/

### Data sets

We used 14 Papuans, 71 WestEurasians, 72 East Asians and 39 South Asians individuals from the Simons Genome Diversity Project (SGDP) (Mallick *et al*. 2016), 40 Papuans from (Malaspinas *et al*. 2016) and an additional 35 Papuans (Vernot *et al*. 2016).

### Filtering variants in real data

We used two sets of outgroups. One is all Sub-Saharan Africans (populations: YRI, MSL, ESN) from the 1000 Genomes Project (Genomes PROJECT *et al*. 2015) and all Sub-Saharan African populations from SGDP (Mallick *et al*. 2016) except Masai, Somali, Sharawi and Mozabite, which show signs of out-of-Africa admixture. The other outgroup is all individuals from the 1000 Genomes Project (Genomes Project *et al*. 2015) plus all non-Papuans from SGDP. For all human data sets, we also removed sites that fell within repeatmasked (Smit *et al*. 2013) regions, and sites that were not in the strict callability mask for the 1000 Genomes Project.

#### Repeat mask regions

hgdownload.cse.ucsc.edu/goldenpath/hg19/bigZips/chromFaMasked.tar.gz Strict callability mask for 1000 genomes: ftp://ftp.1000genomes.ebi.ac.uk/vol1/ftp/release/20130502/supporting/accessible_genome_masks/StrictMask/

The background mutation rate was calculated using the variants density of all variants from populations YRI, LWK, GWD, MSL and ESN in windows of 100 Kb divided by the mean variant density of the whole genome.

### Comparison to Sstar and Conditional Random Field

We called Neanderthal and Denisova segments in the 14 Papuans and compared them to the segments called with CRF with more than 50 posterior probability (Sankararaman *et al*. 2016) available at: https://sriramlab.cass.idre.ucla.edu/public/sankararaman.curbio.2016/ The path to the haplotypes is: summaries/2/denisova/oceania/summaries/haplotypes/CRHOM.thresh-50.length-0.00.haplotypes

We called Neanderthal and Denisova segments in the 35 Papuans and compared them to the segments called with Sstar with more than 99 posterior probability (Vernot *et al*. 2016) available at: https://drive.google.com/drive/folders/0B9Pc7_zItMCVWUp6bWtXc2xJVkk The path to the haplotypes is: introgressed_haplotypes/LL.callsetPNG.mr_0.99.den_calls_by_hap.bed.merged.by_chr.bed

## Acknowledgments

RD was supported by Wellcome Trust grants WT206194 and RG89781. LS was supported by grants 1323-00076 and 6108-00385 from the Danish Council for Independent Research, Natural Sciences (To MHS).

## Supporting Figure legends

**Supporting figure 1 – Demographic parameters for simulation**. The effective population sizes, split times and bottleneck population sizes are shown for the simulated populations.

**Supporting figure 2 - Total segments and sequence called SIM**. The first column show the total number of segments found and the second column show the total amount of sequence that these segments add up to. The rows are different simulation scenarios and the colors of the stacked bar plot show the amount/number of segments that are not found using posterior decoding, where less than half of the segment overlap with the true archaic segments or where more than half of the segment overlaps with the true archaic segment.

**Supporting figure 3 – Effect of adjusting cutoff for when to include a putative archaic segment**. The rows are different simulation scenarios and the columns are different classifications of putative archaic segments. False is segments with zero overlap to the true archaic segments, found<50% are archaic segments that overlap with less than 50% with the true archaic segments and found>50% are segments that overlap with more than 50% with the true archaic segments. On the x-axis is the mean posterior probability of an archaic segment and the y-axis is the amount of sequence left when applying the filter as a fraction of that found with a filter value of 50%.

**Supporting figure 4 - Parameter estimation of Papuans**. The different subpanels show the estimates for the parameters t_admix, a, T_ingroup and T_archaic depending on which outgroup was used (Sub-Saharan Africans) or the whole world (non-Papuans). There is a separate bar for each individual, and the bars are colored according to which dataset they came from.

**Supporting figure 5 - Segment distributions as a function of posterior probability**. Distributions of the number (left) and total length (right) of segments with mean posterior probability as on the x axis. Numbers are given for all 87 Papuans, called with a threshold of 0.5.

**Supporting figure 6 - Length distribution of inferred segments for other methods**. The length distribution of all Denisova and Neandertal segments found using conditional random field (CRF), the hidden Markov model (HMM) and Sstar. For our HMM, Neanderthal are those segments that are shared with other non-African populations and Denisova are those unique to Papuans.

**Supporting figure 7 - Length distribution of Asians, Europeans and Papuans**. The length distributions of segments unique to Papuans (Denisova) and segments shared with other non-African populations (Neanderthal) are shown for segments found using four different population groups.

