## Supplementary Materials for "Detecting archaic introgression without archaic reference genomes"

### Transition matrix

Ingroup to Ingroup and Ingroup to Archaic

#### Haploid case

| Ingroup | Ingroup |
|---|---|

$$P(I \to I) = e^{-T_{admix} \cdot r \cdot L} + (1 - e^{-T_{admix} \cdot r \cdot L})(1 - a)$$

No recombinaiton

Recombination with another Ingroup segment

When $T_{admix} \cdot r \cdot L$ is small we can approximate

$$e^{-T_{admix} \cdot r \cdot L} \approx 1 - T_{admix} \cdot r \cdot L$$

$$\boxed{P(I \to I) = 1 - a \cdot T_{admix} \cdot r \cdot L}$$

The probability of changing state to the archaic state is then:

$$\boxed{P(I \to A) = a \cdot T_{admix} \cdot r \cdot L}$$

#### Diploid case

| Ingroup | Ingroup |
|---|---|
| Ingroup | Ingroup |

$$P(I \to I) = e^{-T_{admix} \cdot r \cdot L} + (1 - e^{-T_{admix} \cdot r \cdot L})(1 - 2a)$$

No recombinaiton

Recombination with another human segment

The 2a is because we dont want to recombine with archaic segment ot either chromosome. The rest of the calculations are the same.

$$\boxed{P(I \to I) = 1 - 2a \cdot T_{admix} \cdot r \cdot L}$$

$$\boxed{P(I \to A) = 2a \cdot T_{admix} \cdot r \cdot L}$$

### Transition matrix

Archaic to archaic and archaic to Ingroup

#### Haploid case

| | |
|---|---|
| Archaic | Archaic |

$$P(A \to A) = e^{-T_{admix} \cdot r \cdot L} + (1 - e^{-T_{admix} \cdot r \cdot L})a$$

No recombinaiton

Recombination with another archaic segment

$$P(A \to A) = 1 - (1 - a) \cdot T_{admix} \cdot r \cdot L$$

$$P(A \to I) = (1 - a) \cdot T_{admix} \cdot r \cdot L$$

#### Diploid case

| | |
|---|---|
| Archaic | Archaic |
| Ingroup | Ingroup |

$$P(A \to A) = e^{-T_{admix} \cdot r \cdot L} + (1 - e^{-T_{admix} \cdot r \cdot L})a$$

No recombinaiton

Recombination with another human segment

If we assume that Archaic segments are only heterozygous the transition probabilities does not change.

$$P(A \to A) = 1 - (1 - a) \cdot T_{admix} \cdot r \cdot L$$

$$P(A \to I) = (1 - a) \cdot T_{admix} \cdot r \cdot L$$

### Emission values

Ingroup and Archaic

#### Haploid case

Ingroup state

$$\lambda_{Ingroup} = \mu \cdot L \cdot T_{ingroup}$$

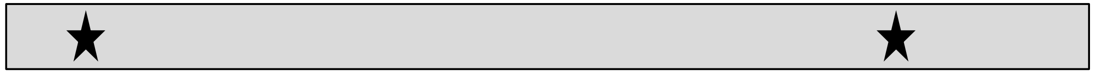

Archaic state

$$\lambda_{Archaic} = \mu \cdot L \cdot T_{Archaic}$$

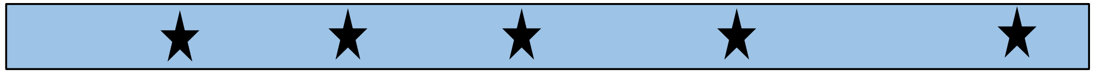

#### Diploid case

Ingroup state

$$\lambda_{Ingroup} = 2 \cdot \mu \cdot L \cdot T_{ingroup}$$

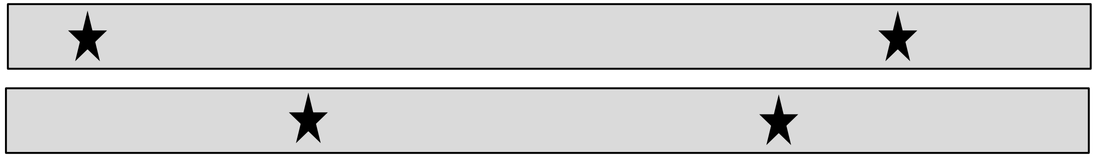

Archaic state

$$\lambda_{Archaic} = \mu \cdot L \cdot T_{Archaic} + \mu \cdot L \cdot T_{ingroup}$$

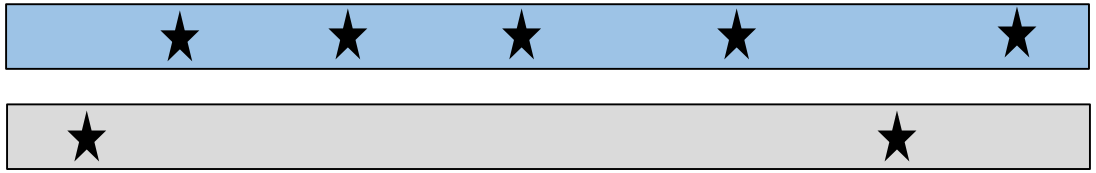

We assume archaic segments are always heterozygous
